# The absence of a neurogenic response to a repeated concussive-like injury in mice

**DOI:** 10.1101/2023.01.16.524157

**Authors:** A. Weingarten, T. M. Madjou, S.N. Yeturu, N. Samudrala, L.E. Villasana

**Author notes:** Corresponding author, Laura Villasana.

## Abstract

In response to traumatic brain injury (TBI), the brain increases its generation of new neurons (neurogenesis) within the hippocampus, a brain region critical for learning and memory. Because neurogenesis plays important roles in learning and memory, post-traumatic neurogenesis may represent an adaptive response contributing to cognitive recovery. In contrast to increases in neurogenesis acutely after injury, levels of neurogenesis become impaired long after TBI. And although chronic deficits in neurogenesis after TBI have been reported by multiple groups, it is unknown whether the hippocampus remains capable of eliciting another neurogenic response to a repeated injury. To address this lack of knowledge, we used a closed head injury model that reflects a concussive-like injury or a mild TBI (mTBI) and assessed levels of neurogenesis in male and female adult mice. Mice received one or two mTBI or sham treatments 3 weeks apart. Compared to mice with a single mTBI, proliferation and neurogenesis were blunted in mice that received a second mTBI. This impaired response was unlikely due to a short recovery time between the two mTBIs as the proliferative response to a second mTBI was also impaired when two months were allowed between injuries. We further found that proliferation was impaired in the radial-glia like cells despite an intact pool. The mice that received two mTBIs also had a blunted intensity in their GFAP staining. In contrast to reports of aberrant post-TBI neurogenesis, we found that the neurons born after mTBI had normal dendritic branches. Lastly, we found that impairments in the inability to mount a neurogenic response after a second mTBI were associated with deficits in neurogenesis-strategy flexibility in the reversal water maze task. Our data suggests that a loss in the neurogenic response could in part contribute to worse cognitive recovery after a repeated concussion. These data may expose a novel target to help improve long-term cognitive outcome following repeated brain injury.

## Introduction

Mild traumatic brain injury (mTBI) is the most common type of TBI, making up 70-90% of TBIs^1,2^. The majority of cognitive complications from mTBI resolve within weeks to months but some mTBIs, especially when repeated cause long-term complications, including changes in mood stability and memory function^3–7^. Currently, there are no FDA-approved therapies to treat chronic cognitive problems from mTBI. However, the brain is endowed with the ability to robustly increase the generation new neurons (neurogenesis) in response to brain injury^8–10^, which may reflect an endogenous repair mechanism.

Because neurogenesis occurs within the hippocampus, a brain region critical for learning and memory^11–13^, there is wide interests in augmenting post-traumatic neurogenesis to facilitate recovery^14–17^. Moreover, neurogenesis influences a wide array of cognitive processes, many of which are altered by TBI. For example, although neurogenesis was first characterized for its importance in pattern separation^18,19^-the ability to discern overlapping contexts-we have since learned that neurogenesis modulates the stress response^20,21^; is critical for the efficacy of a major class of antidepressants^22,23^; facilitates forgetting^24,25^ and is involved in the long-term consolidation of memories^26–28^; and are now discovering its involvement in the architecture of sleep^29^.. Thus, given its various roles in cognitive function, neurogenesis holds strong therapeutic promise for recovery following head injury.

A critical consideration in the therapeutic utility of post-traumatic neurogenesis is that it does not occur without an expense: Levels of neurogenesis become depleted in the months following moderate to severe TBI^30,31^. While these deficits themselves may point to progressive cognitive pathologies associated with neurogenesis-sensitive cognitive domains, loss of an endogenous repair mechanism to guard against subsequent head injuries may also present a critical complication to recovery from repeated concussions. In this study, we used a mild TBI mouse model of a concussion to address whether the hippocampus is capable of increasing the generation of new neurons in response to a repeated mTBI and propose that deficits in the neurogenic response could in part explain why repeated TBIs lead to worse cognitive outcomes compared to a single TBI^32–34^.

The mechanisms underlying TBI-induced deficits in neurogenesis are not clear but may involve changes in neural stem cells. Under physiological conditions, the normally quiescent radial glia-like cells (RGCs) divide to make copies of themselves (symmetric division), new astrocytes and intermediate progenitor cells (IPCs) (for review see^35^). During their transition to neuroblasts, many of the IPCs become phagocytosed by microglia while the remaining surviving IPCs take on a neuronal fate and become immature granule cells within the dentate gyrus. Although changes in proliferation, differentiation and/or survival could contribute to deficits in neurogenesis after TBI, Neuberger et al.,^30^ observed long-term impairments in proliferation following a lateral fluid percussive injury. Because increases in neurogenesis are dependent on the proliferation of RGCs^36^, here we assess whether the proliferative response of RGCs to a subsequent mTBI is altered.

The functional significance of post-TBI neurogenesis is not clear and while it is clear that neurons born after TBI survive and functionally integrate within the hippocampal circuitry^37,38^, their contribution (beneficial or harmful) to recovery is debatable. While several studies suggests that post-traumatic neurogenesis is beneficial for cognitive recovery^39–41^, more recent studies question this type of contribution and instead suggests that reducing new neurons after brain injury may be more beneficial for recovery^31,42,43^. Indeed, new neurons are important for various aspects of cognitive function however, several reports demonstrate that post-TBI born neurons are not the same as those generated in a non-injured brain. For example, neurons generated after controlled cortical impact (CCI) injury mislocalize within the hippocampal circuitry and have aberrant dendritic morphologies^38,44,45^. Notably, it is not known whether a less severe TBI results in similar developmental abnormalities of post-TBI new neurons. Thus, the discrepancy among studies addressing the functional significance of post-traumatic neurogenesis may be owed to the type or degree of injury. To this end, in this study we used a Doublecortin Cre mouse crossed with a reporter mouse to pulse label and trace the dendrites of new neurons generated after a mTBI. In a separate study, neurogenesis-sensitive strategy flexibility was assessed using the reversal water maze task in C57Bl/6J wild-type mice one month after a single or repeated mTBI.

The results of our study suggest that a history of a mTBI impairs the ability of the hippocampus to increase its generation of new neurons in response to a second mTBI. Additionally, we show that the neurons generated after a mTBI have normal dendritic morphologies and that the inability to increase neurogenesis after a repeated mTBI is associated with impairments in strategy flexibility. Taken together our results shed light on a novel pathology in repeated TBI. The finding that mTBI does not generate aberrant neurogenesis warrants efforts towards interventions to prevent or restore the neurogenic capacity following mTBI in order to determine its contribution to cognitive recovery.

## Methods

### Animals

All procedures were performed according to the National Institutes of Health Guidelines for the Care and Use of Laboratory Animals and were in compliance with approved IACUC protocols at Oregon Health & Science University and Legacy Research Institute. Subjects were two-month-old male and female mice housed in standard cages in a 12h/12h light-dark cycle with free access to food and water. Mice were randomly assigned to four experimental groups: sham-sham, sham-mTBI, mTBI-sham, and mTBI-mTBI. To assess neurogenesis, we used POMC-GFP mice which express GFP in immature neurons within hippocampal dentate gyrus as previously shown^37,46^. C57Bl/6J wild-type male and female mice were used to assess proliferation and for behavior testing in the water maze task. To analyze the dendritic morphology of neurons generated after TBI we used Doublecortin-CreER^T2^ mouse (generously provided by Dr. Zhi-Qi Xiong, Institute of Neuroscience, Shanghai, China) which we crossed with a Rosa26-CAG-tdTomato reporter mouse and referred to here as DcxCre/tdTom mice. The tamoxifen-inducible Cre in this mouse is under the control of the Dcx promoter, allowing us permanently pulse label new neurons as previously shown^37^. Litter mate mice were randomly assigned to four experimental groups: sham-sham, sham-mTBI, mTBI-sham, and mTBI-mTBI. Another cohort of wild-type mice received mTBI in order to compare cell death to tissue to similarly aged wild-type mice that received a CCI injury from a separate study.

### Closed head injury and Neuroseverity scores

Mice were anesthetized in an induction chamber using spontaneously inhaled isoflurane (2%) and transferred to a stereotaxic apparatus. Anesthesia was maintained using a nose cone. A Leica Impact One device was used to position a 3mm stainless steel piston flush with the skull surface, which was then retracted and lowered by 2.2mm. The piston was fired at a speed of 4.67m/s with an 800ms dwell time. Mice were allowed to recover on a heating pad. Sham mice received an equal amount of time under anesthesia with no injury. Thirty minutes after injury, mice were assessed for sensorimotor and locomotor deficits using a neurological severity score (NSS) which included the following assessments: gait, exploratory behavior; string test, beam balance and a startle response. The assessments were rated 0-2 with 2 being grossly impaired by an experimenter blind to the treatment of mice. Mice received a second sham or mTBI treatment three weeks or two months later.

### BrdU/EdU/tamoxifen injections

POMC-GFP mice were injected with the thymidine analogue bromodeoxyuridine (BrdU, i.p. 50mg/kg) twice daily (4 hours apart) for 3 days starting 3 days after the second sham or mTBI treatment. Double labeling of BrdU with the immature marker, doublecortin (Dcx) provided a second measure of neurogenesis in addition to the GFP marker. To label proliferating cells, wild-type mice were injected with ethynyldeoxyuridine (Edu, i.p. 50mg/kg) 3 times daily (2 hours apart) 3 days after the second sham or mTBI treatment and were euthanized the following day. To label the dendrites of new neurons born after sham or mTBI treatment, DcxCre/tdTom mice were injected with tamoxifen (i.p. 40mg/kg) twice daily (7 hours apart) for 3 days starting one week after sham or mTBI treatment. Mice were euthanized 2 months later.

### Immunohistochemistry

Mice were deeply anesthetized, transcardially perfused with 0.9% saline followed by 4% paraformaldehyde according to IACUC-approved methods. Brains were removed, post-fixed overnight, cryoprotected in 30% sucrose and frozen. Three to four coronal sections (70μm thick) of the hippocampus approximately spanning ^-^1.46 - ^-^2.8 were selected from each mouse, washed and permeabilized in 0.5% Triton in PBS (PBST). Sections stained for BrdU were treated with 2N HCl at 37°C for 30 minutes, followed by neutralization in 0.1M sodium tetraborate for 10 minutes and washed with PBST (pH 7.4) for 30 minutes. EdU staining was performed using a Click-iT EdU Imaging Kit (Invitrogen). All sections were then blocked using a buffer of 4% normal donkey serum in PBST, then incubated overnight at 4°C with primary antibodies diluted in 2% normal donkey serum in PBST. Primary antibodies were as follows: rabbit anti-GFP (1:400; Invitrogen); guinea pig anti-Dcx (1:500; Millipore); rabbit anti-Ki67 (1:400; Abcam), rat anti-BrdU (1:500; Santa Cruz), rat anti-Sox2 (1:400; Invitrogen), mouse anti-glial fibrillary acidic protein (GFAP; 1:600; Novus); rabbit anti-GFAP (1:600; Dako); and rabbit anti-tdTom (1:500; Clonetech). Edu was stained according to the manufacturer’s recommendations (Click-iT kit, Thermofisher Scientific). Sections were washed the following day and incubated with secondary antibody solutions for 2 hours at room temperature. Secondary antibodies were as follows: donkey anti-rat (1:500; Alexa Fluor 555, Invitrogen), donkey antirabbit (1:500; Alexa Fluor 488, Invitrogen), goat anti-guinea pig (1:500; Invitrogen); donkey antimouse (1:500; Alexa Fluor 647, Invitrogen). Lastly, sections were washed with PBST, incubated with DAPI in PBST (1:10,000) for 20 minutes, then mounted with Vectashield Anti-Fade Mounting Medium.

Fluoro-Jade C staining was performed on mice sacrificed 3 hours after injury sham or mTBI treatment. Slices (70μm thick) taken from directly beneath the injury site were mounted and dried on Colorfrost Plus slides, ethanol rehydrated, incubated in 0.06% potassium permanganate for 10 minutes followed by 0.002% Fluoro-Jade C (Histo-Chem Inc.) for 10 minutes. Slices were cleared with xylene and coverslipped with castor oil.

### Image Analysis

Slides were coded for blind analyses and imaged using a Leica Mi8 confocal microscope with a 20x/0.8 NA lense. To assess neurogenesis, GFP and Dcx^+^/BrdU^+^ cells within the granular cell layer (GCL) of the dentate gyrus were imaged through a 10μm stack. These included the infra- and suprapyramidal blades of the dentate gyrus. To assess proliferation, Ki67^+^, Edu^+^, and BrdU^+^ cells within the GCL were imaged through a 20μm stack. Radial glia-like cells within the granule cell layer of the dentate gyrus were identified using GFAP/Sox2 labeling and by their primary process. The density of cells was calculated by normalizing the number of cells to the GCL volume (area multiplied by the depth). The percent of proliferating RGCs was calculated by the density of GFAP^+^/Sox2^+^/Ki67^+^ cells divided by the density of GFAP^+^/Sox^+^ cells. To assess the dendrites of new neurons for Sholl analysis (Image J2 version 2.3.0/1.53f), individual tdTom^+^ cells were imaged through their entire dendritic tree. Only tdTom^+^ cells that did not have truncated dendrites due to sectioning were included in the analyses. Because neurogenesis frequently occurs in clusters, 3D reconstructions were made to identify overlapping somas and correctly identify the dendrites from individual cells. For each mouse, 8-10 cells from 3-4 different sections were traced. For intensity analyses of GFAP, z-stacks of 7μm were obtained at consistent acquisition settings and mean intensity of pixels of the dentate gyrus GCL were quantified using ImageJ after digital subtraction of mean background values.

### Rotorod testing

Mice were placed on a rotating rod (Rotomex-5, Columbus Instruments, Columbus, OH) initially rotating at 5 rpm by an experimenter blind to the treatment of the mice. Once all 4 mice were placed on individually divided sections of the rod, the speed of the rod was automatically increased by 1 rpm every 3s for a maximum of 24 rpm. Beam breaks digitally recorded the fall of each mouse. Three trials were conducted 5 minutes apart for each mouse approximately 2 hours after the second sham or mTBI treatment. The rotorod was only conducted on the mice with a two-month injury interval.

### Reversal water maze testing

Four weeks after the second sham or mTBI treatment, mice were tested on the water maze by an experimenter blind to the treatment of the mice. One hour prior to daily morning testing, mice were moved to a holding room separate from the water maze. They were placed in individual cages that sat half way on top of a heating pad. The water maze (122cm circular pool) was placed the middle of a room with stationary distal cues and filled with opaque water (white Crayola color) maintained at 20°C. A clear circular platform (10cm in diameter) invisible to the mice (2cm under the water surface) was located in the middle of one of four imaginary quadrants of the pool. Mice were trained to locate the platform with 4 trials per session (10 min inter-trial-interval) with one session per day for 3 days. Mice that did not locate the platform before 60 seconds were hand-led to the it and all mice were allowed to remain on the platform for 5 seconds. On the 4^th^ day, the platform was removed and the time the mice spent searching in the quadrant where the platform was previously located was assessed as well as the search proximity from the platform location. On the 5^th^ day, the location of the platform was moved to the opposite quadrant and the mice were trained to locate the new location for 3 days followed by a second probe trial on the 8^th^ day. Ethovision (Noldus; Netherlands) was used to analyze the performance of mice.

### Statistical Analysis

All data were first assessed for normality and homogeneity of variance to determine the use of parametric versus non parametric statistical tests as indicated in the Results section. Data are expressed as Means ± SEM and analyzed with SPSS Statistics (IBM, Armonk, NY) and group comparisons were considered significant at p < 0.06. All figures were generated using Prism Software (GraphPad Software, La Jolla CA). There were no sex differences in our analysis of neurogenesis and proliferation in the three-week injury interval experiment, thus the sexes were combined for the experiment involving a 2 month injury interval. A one-way ANOVA was used to analyze group differences in treatment, which consisted of four groups: sham-sham; sham-mTBI; mTBI-sham; and mTBI-mTBI. Where appropriate, Tukey’s post-hoc test was used to assess group differences. Data involving Edu and BrdU quantification were normalized to the control group (sham-sham) within their cohorts. Linear regressions were used to analyze the relationship between neuroseverity scores (NSS) and BrdU cell densities; and BrdU and Ki67 cell densities. The water maze learning curves were analyzed with sessions as the within-subject variables using a repeated-measures ANOVA (RM-ANOVA). Sholl analysis was conducted using a RM-ANOVA with distance from the soma as the within subject variable.

## Results

### mTBI resembles a concussive-like injury

Unlike a more severe controlled cortical impact injury (CCI), we found no evidence of gross cell death hours after injury (Fig. 1A). The time it took mice to rise on all limbs after surgery (righting reflex) was significantly longer in mTBI mice (Fig. 1B; p < 0.0001, Student’s t-test). Neurologic severity scores assessing gait, exploratory behavior, string test, beam balance, and startle response 30 min after surgery showed deficits in mTBI mice (Fig. 1C; p < 0.0001, Student’s t-test). mTBI mice moved less and slower than sham mice in the open field (Fig. 1D, E; p < 0.01, Student’s t-test).

**Fig. 1.**
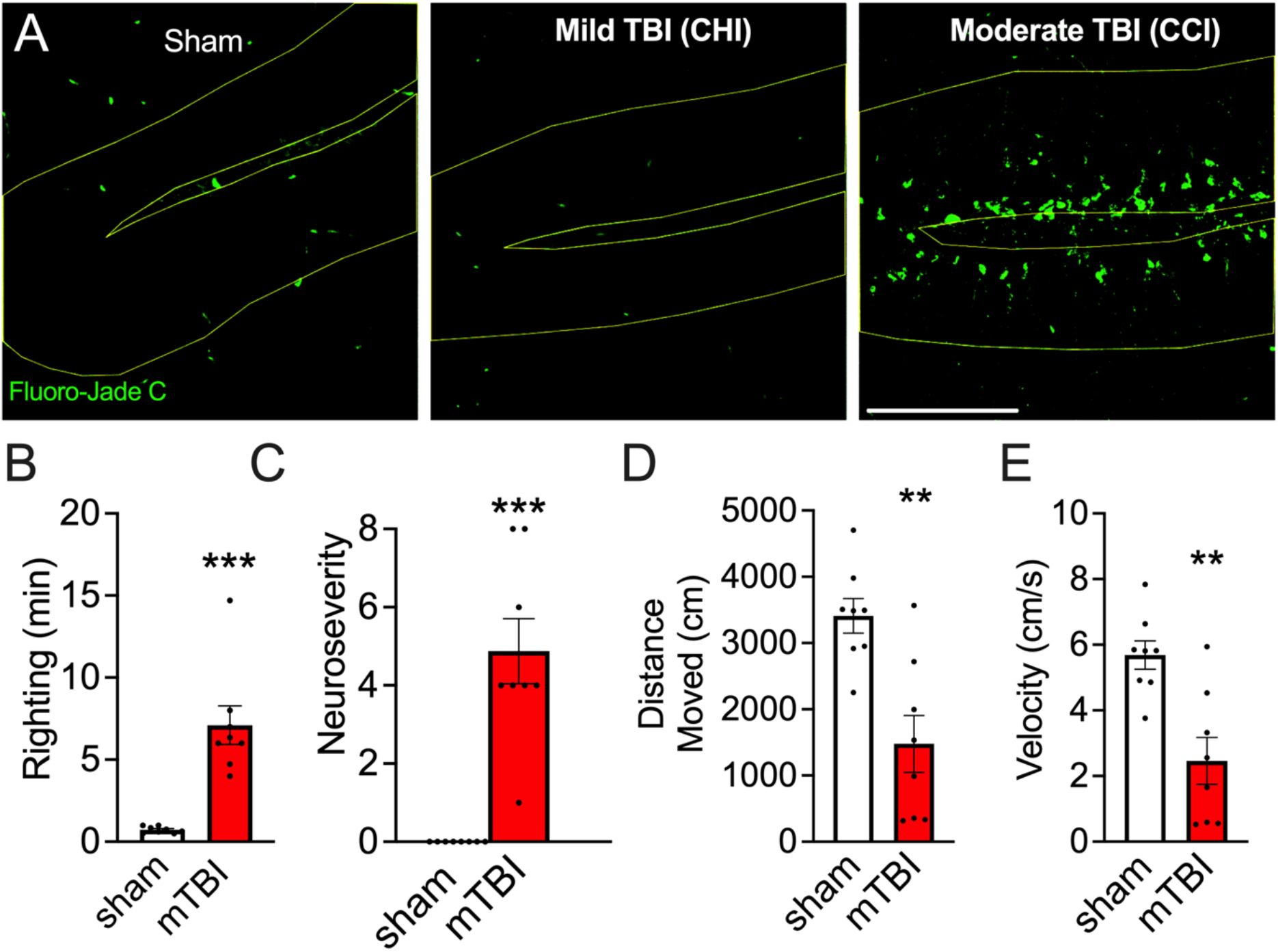
mTBI resembles a mild concussive-like injury. **A**. Cell death assessed in the dentate gyrus 3 hours after a controlled cortical impact (CCI) or mTBI. **B.** Righting reflex of mice immediately following sham or mTBI treatment. **C**. Greater neuroseverity scores 30min after surgery in mice with a mTBI. **D**. mTBI mice moved less and slower **(E)** in the open field 30 min after surgery. *p < 0.05; **p < 0.01; ***p < 0.001. Scale bar 100μm.

### Increases in neurogenesis are absent in after a repeated mTBI

To determine if a second mTBI elicits a neurogenic response, POMC-GFP mice underwent two treatments (sham or mTBI) three weeks apart and were injected with the mitotic marker BrdU 3 days after the second treatment (i.p. 150mg/kg, twice daily 4 hours apart) for 3 days (Fig. 2A). The NSS of mice after the second mTBI were no different than the NSS of mice with a single mTBI (Fig. 2B). There was a significant effect of treatment in the density of GFP^+^ cells (Fig. 2C & D; F_3,20_ = 4.95; p < 0.0001). Mice with a single mTBI had greater densities of GFP^+^ cells compared to sham-sham mice (p < 0.05) and compared to mTBI-mTBI mice (p < 0.05). In contrast, mTBI-mTB mice had similar densities of GFP^+^ cells as sham-sham mice. There was a similar effect of treatment in the density of Dcx^+^ cells (Fig. 2E & F; F_3,20_ = 13.11; p < 0.0001). Mice with a single mTBI had a greater density of Dcx^+^ cells compared to sham-sham mice (p < 0.001), mTBI-sham mice (p < 0.0001) and mTBI-mTBI mice (p < 0.01). The lack of increase in the density of immature neurons following a second injury occurred in the absence of deficits in baseline levels of neurogenesis (Fig.3D & F, green versus gray bars). There were no significant sex differences or sex by treatment interactions in either measures of proliferation.

**Fig. 2.**
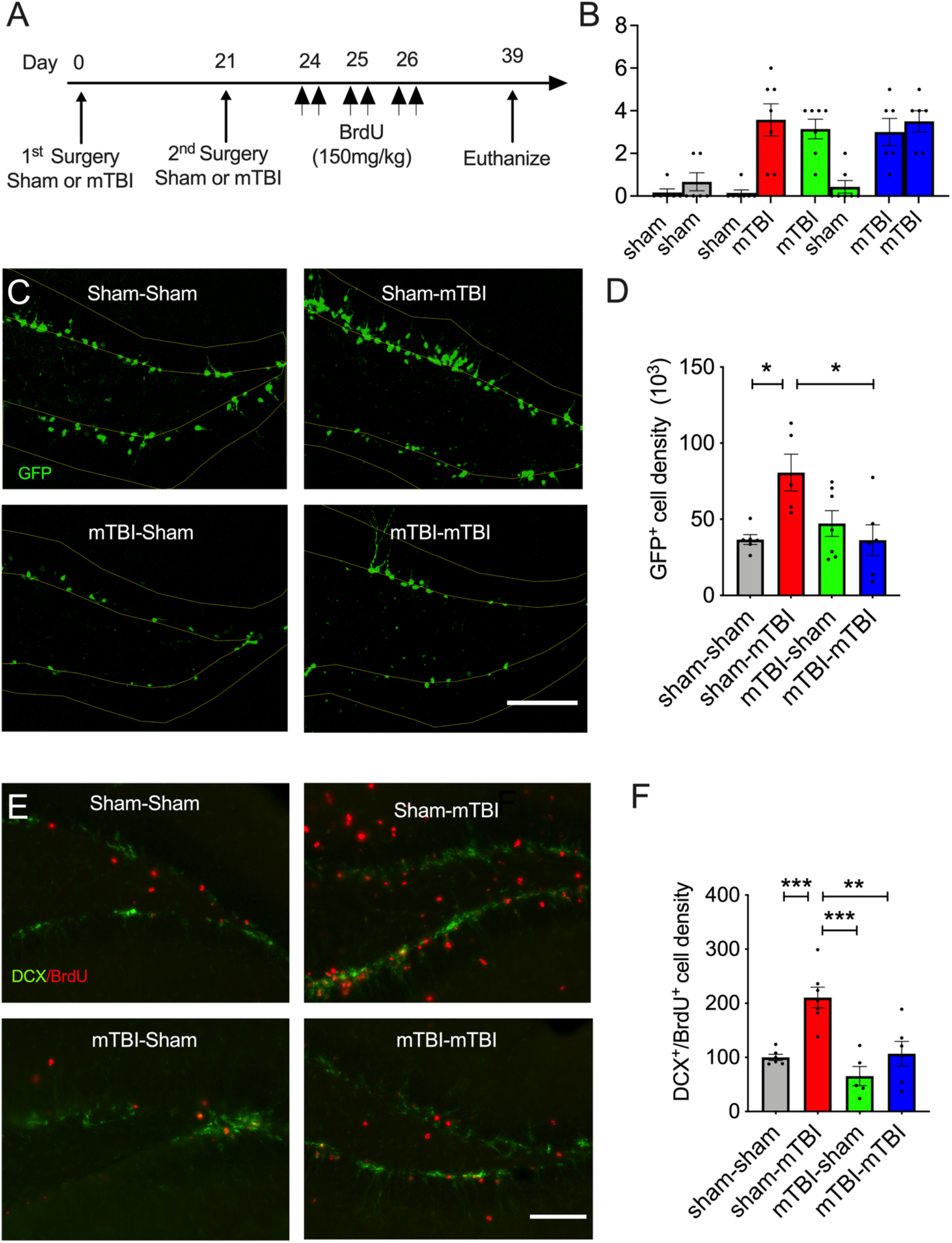
A prior mTBI impairs the neurogenic response to a second mTBI. **A.** Experimental timeline. **B.** Neuroseverity scores (NSS) 30 min after the 1^st^ and 2^nd^ sham or mTBI treatment. **C.** Images of the dentate gyrus stained with GFP and its quantification **(D)** shows increased neurogenesis in mice after one mTBI (red bar) but not after a second mTBI (blue bar). A similar pattern was observed with the immature neuronal marker doublecortin (Dcx) co-stained with BrdU in **E & F**.

**Fig. 3.**
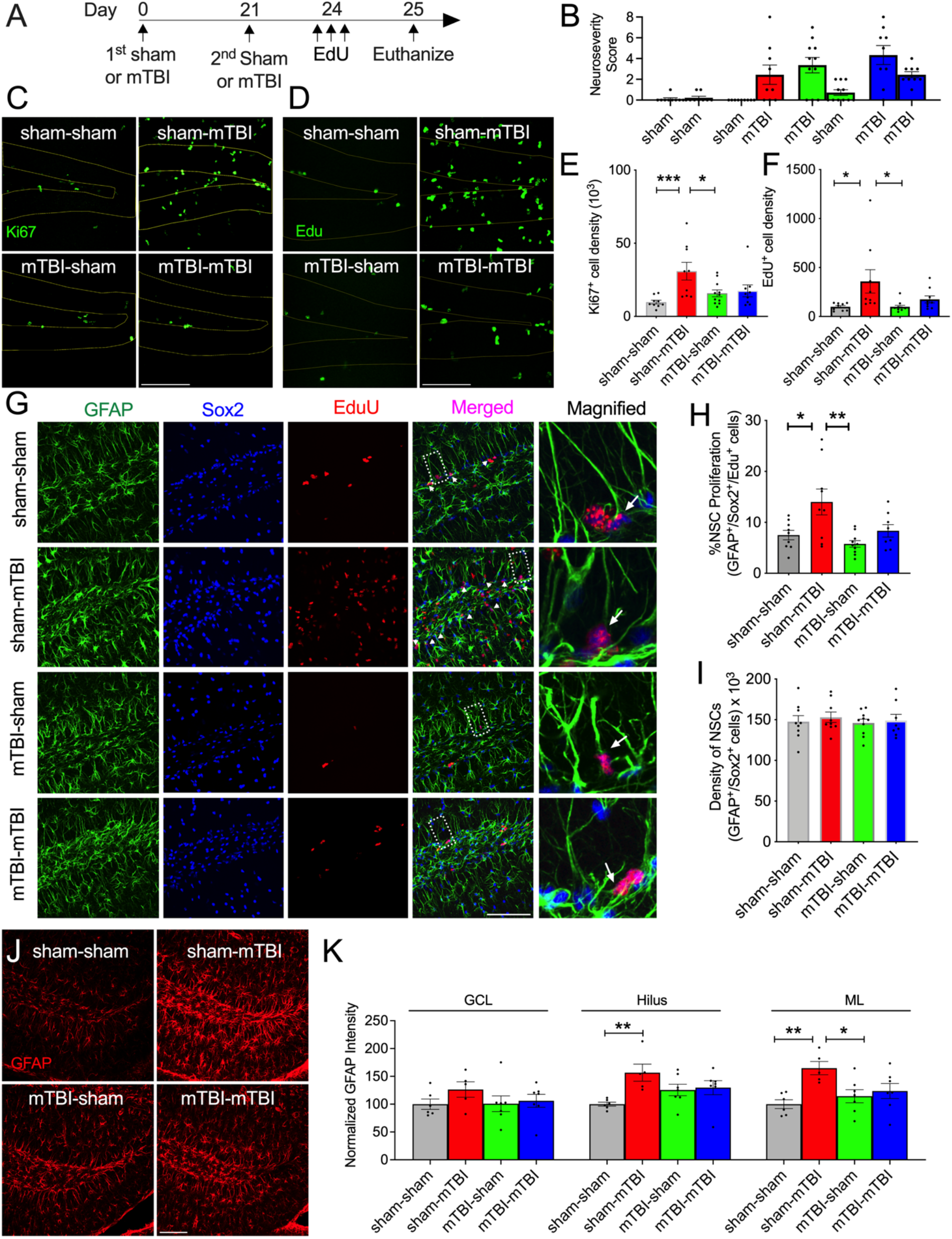
Absent proliferative response of radial glia-like cells to a second mTBI. **A.** Experimental timeline. **B.** NSS 30 min after the 1^st^ & 2^nd^ treatment shows similar degrees of neurologic injury in sham-mTBI and mTBI-mTBI mice (red versus blue bars). **C**. Confocal images of the dentate gyrus (traced in white) stained with the proliferative marker Ki67 and its quantification **(E)** shows increased proliferation after one mTBI (red bar) but not after a second mTBI (blue bar). A similar pattern was observed with an additional proliferative marker EdU **(D, F**). **G.** Confocal images of proliferating RGCs in the dentate gyrus identified with GFAP/Sox2/Edu co-staining. Examples of proliferating RGCs are noted by white arrows in the magnified images (right column). **H.** Increased RGC proliferation in mice after a single mTBI (red bar) but not after a second mTBI (blue bar). **I**. There were no group differences in the density of RGCs (GFAP^+^/Sox^+^ cells). **J**. Representative images of the dentate gyrus stained with the astrocytic marker GFAP and its quantification (**K**). *p < 0.05; **p < 0.01; ***p < 0.001. Scale bar 100μm.

### Impaired proliferative response of RGCs to a subsequent mTBI

To determine whether impairments in the neurogenic response after a repeated mTBI were associated with impairments in proliferation, wild-type mice underwent two treatments (sham or mTBI) three weeks apart and were injected with the mitotic marker EdU 3 days after the second surgery (i.p. 50mg/kg, 3 times every two hours (Fig. 3A). There were no differences in the NSS of sham-mTBI and mTBI-mTBI mice after the second treatment (Fig. 3B; red versus blue bars). There was a treatment effect on the density of Ki67^+^ cells (Fig. 3C & E, p = 0.008, Kruskal Wallis). Mice with a single mTBI had higher densities of Ki67^+^ cells compared to sham-sham treated mice (p < 0.001, Mann-Whitney) and compared to mTBI-sham treated mice (p < 0.05, Mann-Whitney). In contrast mice with a subsequent mTBI did not have significantly different densities than sham-sham mice. A similar effect of treatment was observed in the density of EdU^+^ cells (F_3,34_ = 4.13; p = 0.013). Sham-mTBI treated mice had higher densities of EdU^+^ cells compared to sham-sham (p < 0.05) and mTBI-sham treated mice (p = 0.05). Again, this deficit was observed in the absence of baseline levels of proliferation (green bars). The NSS of mice after a single mTBI and mice with a subsequent mTBI did not differ (Fig. 3B). There were no significant sex differences or sex by treatment interactions.

Increases in neurogenesis after TBI are driven by the proliferation of RGCs^36^. Thus, we next determined whether there were group differences in proliferation specifically within the RGCs. There was a treatment effect on the density of GFAP^+^/Sox2^+^ cells co-labeled with EdU (Fig. 3G & H, p = 0.02, Kruskall-Wallis). Sham-mTBI mice had a greater density of proliferating RGCs than sham-sham (p < 0.05, Mann-Whitney) and mTBI-sham treated mice (p < 0.01, Mann-Whitney). There were no deficits in baseline levels of RGC proliferation (Fig 3H, green bar) and notably, there were no deficits in the density of total RGCs between the different groups (Fig. 3I).

Unsurprisingly the intensity of GFAP staining appeared to differ between groups, thus we quantified the pixel intensity of each group within the different regions of the dentate gyrus. Although the sham-mTBI group appeared to have higher GFAP intensities within the GCL, an effect of treatment did not reach significance (Fig. 3J & K). However, there was a treatment effect in the hilus (F_3,21_ = 3.80; p = 0.03). Sham-mTBI mice had higher GFAP intensity levels compared to sham-sham mice (p < 0.01). There was also a treatment effect within the molecular layer (ML, F_3,21_ = 4.75; p = 0.11). Sham-mTBI mice had greater GFAP intensities than sham-sham (p < 0.01) and mTBI-sham treated mice (p < 0.05).

### Increasing the injury interval did not ameliorate impairments in the proliferative response

We considered the possibility that the RGCs may require a longer interval between injuries to functionally recover, thus we increased the injury interval to two months (Fig. 4A). To determine whether our results were replicable using different markers of proliferation, we used Ki67 which is expressed in all phases of the cell cycle except the G0 phase, and BrdU which is more commonly used than EdU. Similar to the three-week injury interval, there was an effect of treatment on the density of Ki67^+^ cells (Fig. 4D & E; F_3, 16_ = 30.09; p < 0.01). Sham-mTBI treated mice had increased densities of Ki67^+^ cells compared all other three groups of mice (p < 0.0001). A similar effect of treatment was observed in the density of BrdU^+^ cells (Fig. 4E & G, F_3,16_ = 18.45; p < 0.0001). The density of BrdU^+^ cells was greater in sham-mTBI mice compared to all other three groups (p < 0.0001 versus sham-sham and mTBI-mTBI; p < 0.001 versus mTBI-sham).

**Fig. 4.**
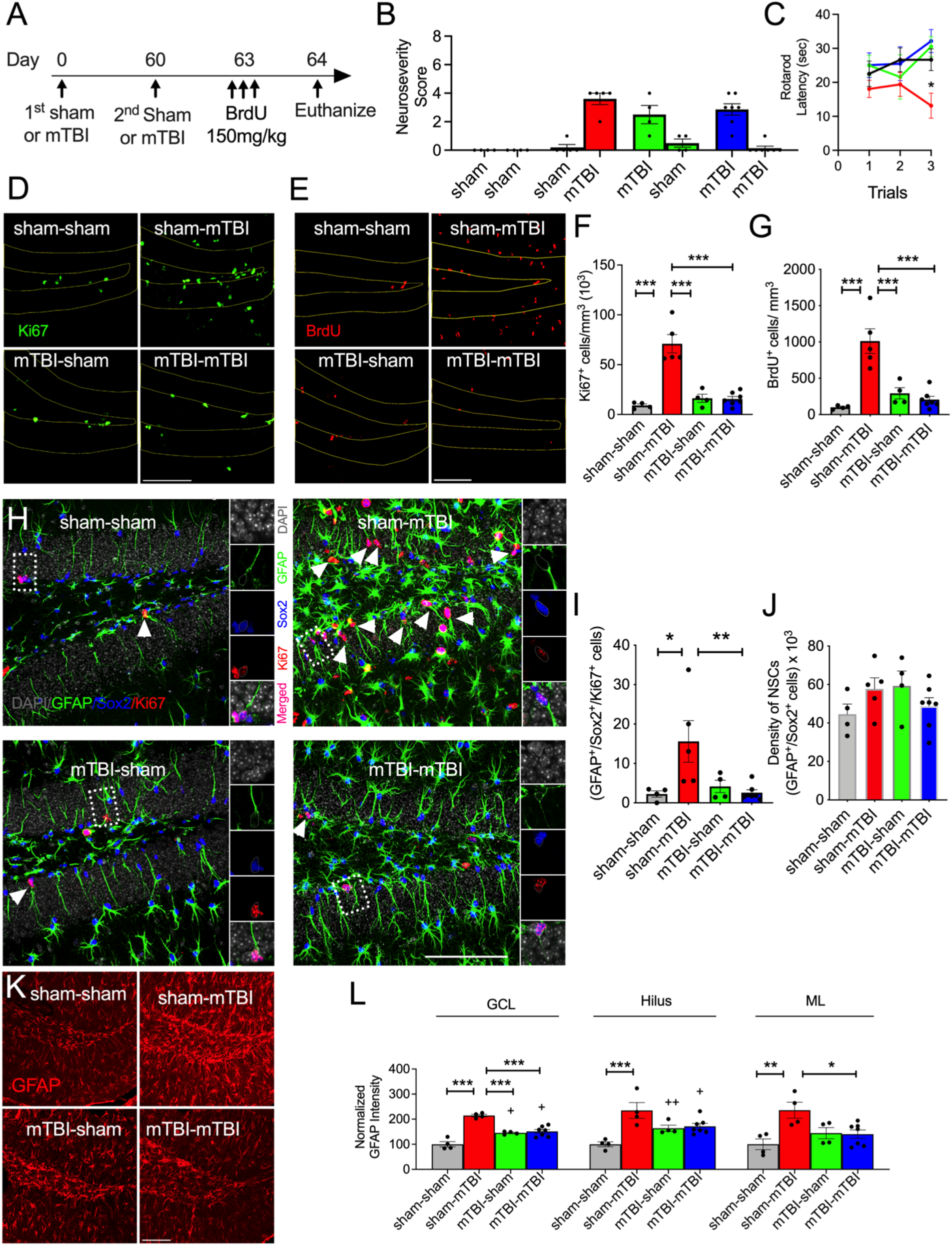
Increasing the injury interval does not ameliorate deficits in the proliferative response of RGCs to a second mTBI. A. Experimental timeline. **B**. Mice with a second mTBI did not demonstrate higher neurologic injury or impaired rotorod performance whereas sham-mTBI mice did. **D**. Images of the dentate gyrus stained with Ki67 and its quantification **(F)** shows increased proliferation after one mTBI (red bar) but not after a second mTBI (blue bar). A similar pattern was observed with an additional proliferative marker BrdU **(E & G**). **H** Confocal images of the dentate gyrus stained with GFAP, Sox2 and Ki67 to identify proliferating RGCs (denoted by arrow heads). Dashed boxes denote magnified examples (right column). **I.** Quantification of proliferating RGCs shows enhanced proliferation of RGCs in the sham-mTBI group but not the group with a second mTBI (red versus blue bars). **J**. Quantification of the density of RGCs shows no group differences. **K.** Representative images of the dentate gyrus stained with the GFAP and its quantification shows the highest levels of GFAP intensity occurred in the sham-mTBI treated mice. p < 0.05; **p < 0.01; ***p < 0.001; ^+^p < 0.05; ^++^p < 0.01. Scale bar 100μm.

Impairments in the proliferative response of RGCs to a repeated mTBI with a two-month injury interval were observed (Fig. 4J & I; p < 0.05, Kruskal Wallis). Sham-mTBI treated mice had greater densities of proliferating RGCs compared to sham-sham and mTBI-mTBI treated mice (p < 0.05 and p < 0.01 respectively). Impairments in the proliferation of RGCs within the mTBI-mTBI group occurred despite a lack of deficits in the RGC pool (Fig. 4J).

A similar group pattern in the intensity of GFAP expression within the different regions of the dentate gyrus was again observed (Fig. 4K-L). There was an effect of treatment in the GCL where the RGCs reside (F_3,15_ = 31.87; P < 0.0001). Sham-mTBI treated mice had greater intensities compared to all other groups of mice (p < 0.0001 versus sham-sham and mTBI-mTBI; p < 0.001 versus mTBI-sham). However, mTBI-sham and mTBI-mTBI mice also had greater intensities than sham-sham mice (p < 0.01). There was also a treatment effect on GFAP intensity within the hilus (F_3, 15_ = 8.23; p < 0.001). Sham-mTBI mice had elevated levels of GFAP staining compared to sham-sham treated mice (p < 0.001). mTBI-mTBI also had greater levels of staining compared to sham-sham treated mice (p < 0.05). Lastly, an effect of treatment on GFAP intensity was also observed in the ML (F_3,15_ = 5.69; p < 0.01). Sham-mTBI treated mice had higher intensities compared to sham-sham (p < 0.01) and mTBI-mTBI treated mice (p < 0.05).

In stark contrast to the 3-week injury interval experiment, we observed that mice with a second mTBI demonstrated strikingly low NSS scores (Fig. 4B). Surprised by this result, we conducted a rotorod test 2 hours after the second sham or mTBI treatment. Although a two-way repeated measures ANOVA did not reach significance (p = 0.08, treatment by trial interaction), analysis of the final trial showed that the sham-mTBI had shorter latencies compared to sham-sham group whereas mTBI-mTBI had similar latencies to the sham-sham group (p < 0.05).

### The density of BrdU^+^ cells after the first mTBI predicts the proliferative response to second mTBI

Previous data showed that the degree of neurologic injury predicts increases in post-TBI neurogenesis^10^. Therefore, we asked whether the degree of proliferation after the first injury also predicted the proliferative response to the second injury. We also observed that the NSS of mice predicted the degree of proliferation after mTBI (Fig. 5C; r square = 0.67; p < 0.001, linear regression). We further observed that the density of BrdU^+^ cells which were labeled 3 days after the first injury, was inversely correlated with the density of Ki67^+^ cells present 3 days after the second injury (Fig. 5D; r square= 0.55; p < 0.05 linear regression), suggesting that impairments in the proliferative response to the second mTBI can be predicted by the degree of proliferative response to the first injury.

**Figure 5.**
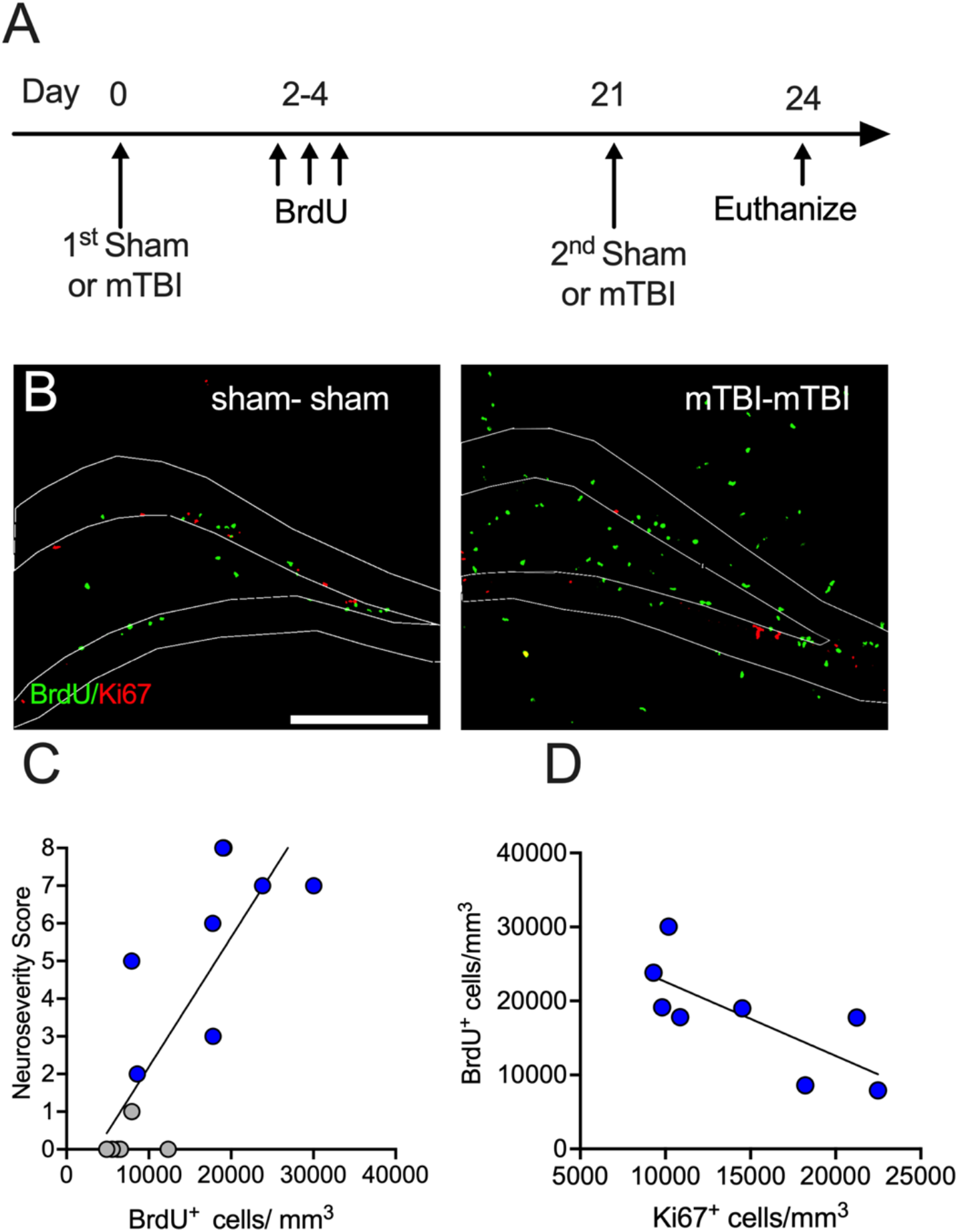
The proliferative response to an initial mTBI predicts the response to second mTBI. **A.** Experimental time line. **B**. Confocal images of the dentate gyrus costained with BrdU and Ki67. **C**. NSS predicts proliferation after an initial mTBI (r square = 0.67; p < 0.001). **D.** Inverse correlation between the degree of proliferation after the first mTBI (BrdU) and the second mTBI (Ki67, r^2^ = 0.55; p < 0.05 linear regression). Scale bar 200μm

### Dendrites of neurons born after mTBI are morphologically typical

Prior studies demonstrate that neurons born after CCI as well as other types of brain insults such as ischemia have aberrant dendritic morphologies^37,38,42^. To determine whether a mild TBI resembling a concussion also results in aberrant neurogenesis, tamoxifen was used to label the new neurons in DcxCre/tdTom mice. The dendritic analysis of 6-8 week old tdTom labeled neurons that were born after a single mTBI demonstrates they are morphologically similar to those born in noninjured mice (Fig. 6A-C). Sholl analysis and dendritic length (Fig. 6A-B) showed very similar dendritic measures between the two groups which is also reflected in the examples of the dendritic traces (Fig. 6C). There was no significant treatment x distance interaction nor a treatment effect on the dendritic length.

**Fig. 6.**
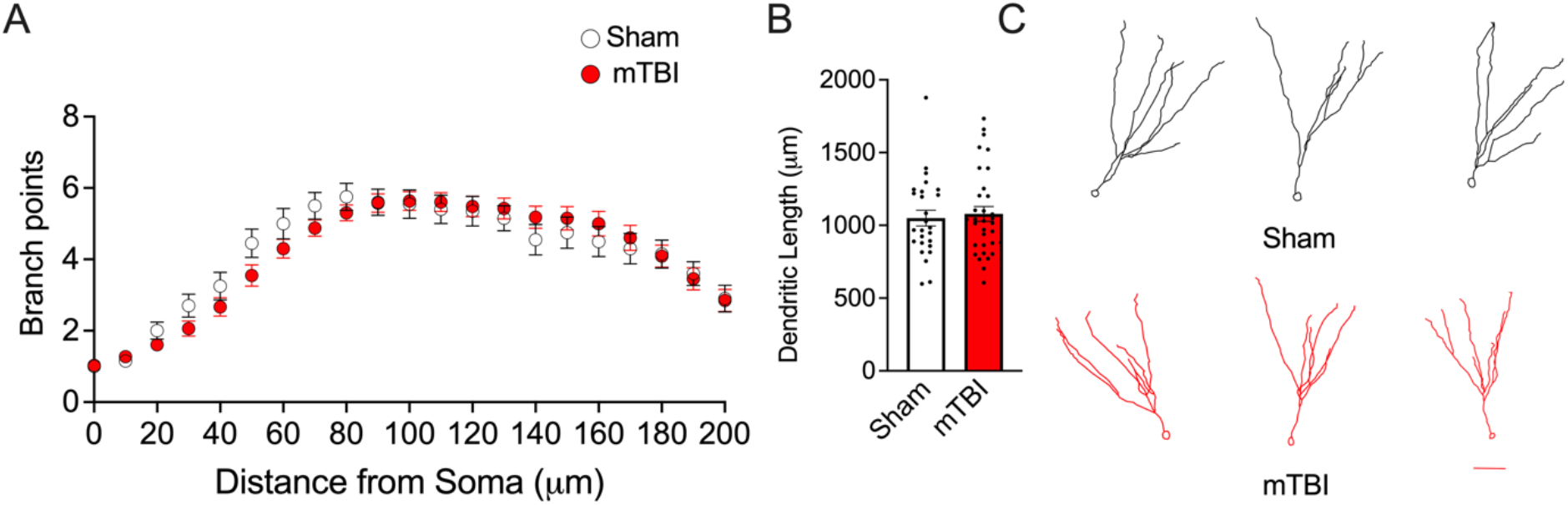
Dendrites of neurons born after mTBI appear typical. **A**. Sholl analysis demonstrates similar branching and length **(B)** of dendrites of new neurons from sham or mTBI treated mice **C.** Dendritic traces of neurons born after mTBI. Scale bar 50μm.

### Impairments in strategy flexibility but not spatial reference memory in mice with repeated mTBI

Mice were tested in the water maze one month after the second sham or mTBI to allow the new neurons to mature and integrate within the hippocampal circuitry. Deficits in neurogenesis do not consistently result in impairments in the reference memory paradigm of the water maze whereas multiple studies have shown that the reversal water maze task, which involves distinguishing overlapping cues (pattern separation), is sensitive to changes in neurogenesis^47–50^. Given the notion that post-TBI increases in neurogenesis facilitates cognitive recovery, the performance of mice on the standard reference versus the reversal water maze test was assessed to determine if a loss in the integration of a robust population of new neurons that are normally generated after injury was associated with worse recovery.

There were no group differences in spatial memory acquisition when mice were trained one month after the last sham or mTBI treatment (Fig. 7A). All mice learned the location of the platform and there were no group differences in swim speed. All groups showed a bias towards the quadrant where the platform was previously located (p < 0.01 sham-sham; p < 0.0001 sham-TBI and mTBI-mTBI), however a few mice within each group did not show this preference (sham-sham, 3; sham-mTBI, 1; mTBI-mTBI, 2). As different weights for the memory of the old platform could potentially influence acquisition to a new platform in the reversal task, the mice that did not show a bias for the new platform during the first probe trial were omitted from the reversal water maze analysis. Fig 7C shows that this exclusion criteria did not affect the results of the reference memory for the first platform.

**Fig. 7.**
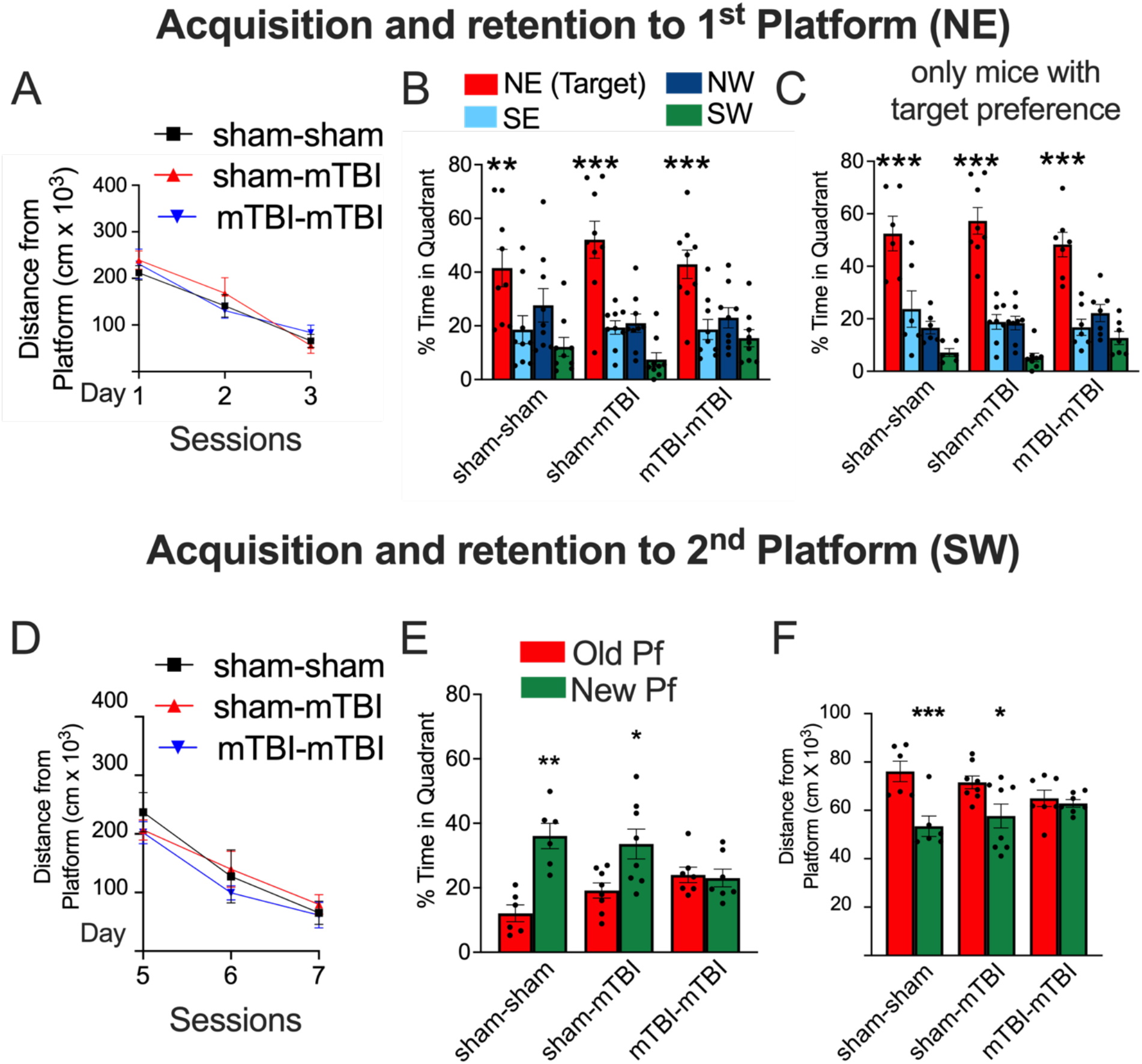
Impairments in strategy flexibility in the reversal water maze in mice with a repeated mTBI. **A.** Spatial acquisition trials to the first platform location. **B.** During the first probe trial, all groups spent significantly more time in the quadrant containing the first platform location. **C.** Only mice that showed a bias for the target quadrant were included for reversal water maze learning and memory. **D**. All mice learned the location of the new platform during reversal training. **E-F.** Sham mice and mice with a single mTBI spent more time in the correct quadrant (E) and searched closer to the correct platform location (F) during the second probe trial, whereas mice with a repeated mTBI did not (*p < 0.05; ** p < 0.01; ***p < 0.001 new versus old quadrant (E) or old platform location (F).

During the training trials to the new platform, there was a subtle trend for the mice with the repeated mTBI to perform worse, however there were no significant differences in group performances (Fig. 7D). Moreover, group performances by the 3^rd^ session of the reversal task were indistinguishable from each other. Analysis of the percent time in the correct quadrant during the first probe test (Fig. 7E) however showed that sham-sham and sham-mTBI mice spent more time in the correct quadrant versus the old quadrant (p < 0.001, p < 0.05 respectively) whereas mTBI-mTBI mice spent a similar amount of time in both quadrants. We also compared the bias to the old platform versus the new platform location using the search proximity to where the platforms where previously located, and found similar results: Sham-sham and sham-mTBI treated mice searched significantly closer to the correct platform location (Fig. 7F; p < 0.01, p < 0.05 respectively) whereas mice with a repeated mTBI did not search closer to the new platform versus the old platform. There were no sex differences in any of the water maze measures.

## Discussion

Acute increases in neurogenesis after TBI are well established. More recent studies however show that constitutive levels of neurogenesis decline in the months following TBI which may be related to changes in the behavior of the neural stem cells^30^. Based on these observations, we asked whether the hippocampus remains capable of eliciting a neurogenic response after a repeated mild TBI. Given that concussions are the most common type of TBI sustained by humans, we explored this question using a mild TBI mouse model. While an initial mTBI increased neurogenesis, not unlike in previous reports, we found that the neurogenic response was absent in mice with a prior mTBI. These observations were noted using the POMC-GFP mouse, an established mouse model for neurogenesis and further corroborated with a doublecortin antibody, a standard marker for immature neurons.

Impairments in the neurogenic response could stem from changes in any of the stages of neurogenesis. In this study, we began interrogating whether deficits in the neurogenic response occurred in conjunction with changes in the proliferative response, the first stage of neurogenesis. Using two different markers of proliferation (Ki67 and BrdU), we found that the proliferative response was likewise blunted in mice with a prior mTBI. We first observed this impairment in mice that had a 3-week interval between injuries. At this short interval, proliferating cells may have still been undergoing higher rates of proliferation or may not have been in a “active” state to induce proliferation^51^. Therefore, we asked if a 2-month interval duration between injuries would result in similar deficits. Again, using two different markers of proliferation, Ki67 and BrdU, the most commonly used thymidine analogue to assess proliferation, we found a similar lack in the proliferative response to a repeated mTBI when the recovery time between injuries was considerably increased. Thus, deficits in the proliferative response were unlikely related to the recovery time required for proliferation to return to baseline levels. Further supporting this conclusion, we did not observe increased proliferation in mice with a single mTBI (mTBI-sham group) at 3 weeks nor at 2 months after injury, demonstrating proliferation had returned to baseline levels.

Although the extent to which a history of a mTBI impairs the neurogenic response to a repeated mTBI is novel, work by Santhakumar’s group shows that a fluid percussion injury (FPI) impairs the neurogenic response to subconvulsive doses of kainic acid^30^. Specifically, they demonstrated that the density and proliferative rates of the intermediate progenitor cells (IPCs) at baseline (i.e. no subsequent challenge) are suppressed 3 months after FPI. Because radial glia-like cells (RGCs) are required for the neurogenic response to TBI, here we focused our examination RGCs and on changes in their proliferative behavior in response to a subsequent challenge (a second mTBI). We found impairments in the proliferative response of the RGCs to a second mTBI in both mice that had a 3-week or a 2-month interval between injuries. Notably, impairments in the proliferative response of RGCs to a second mTBI were observed without deficits in their population, suggesting that impairments in the proliferative behavior of RGCs caused by mTBI occur prior to pool depletion. Furthermore, these findings show that failure in the proliferative response of RGCs to a repeated mTBI are associated with deficits in the neurogenic response. We did not examine whether IPCs undergo similar changes or whether an initial mTBI also alters cell fate or survival. Changes in the latter stages of neurogenesis following an initial mTBI could also alter the neurogenic response to a repeated mTBI.

In agreement with prior studies demonstrating chronic elevations in GFAP after TBI^10,52,53^, sham-mTBI mice had higher intensity levels in GFAP than non-injured mice. Surprisingly however, mice that received a second mTBI had lower GFAP intensity levels than mice that received a single mTBI at the same time point. The dampened elevation in the reactive astrocytic marker after the second injury is reminiscent of delayed preconditioning. For instance, cellular stress prior to a TBI can reduce pathology and neurologic outcome^54^. The surprising observation that mice with a repeated mTBI had no evidence of gross neurologic impairment or a deficit in the rotrorod test, could potentially be explained by a form of delayed preconditioning which was more likely with the 2 month versus the 3 week injury interval. Based on our observations, we speculate that a prior mTBI may act as a preconditioned stimulus to alter the subsequent induction of neurogenesis. As such, we cannot exclude the possibility that impairments in the neurogenic response after a concussive injury may in part be due to a preconditioning effect or increased tolerance rather than a specific dysfunction in the proliferative capacity of the RGCs. It bears noting that preconditioning is typically demonstrated on the scale of hours to days. To the best of our knowledge, the effects of preconditioning on TBI responses have not been demonstrated beyond one week^55^. Further, in the majority of TBI studies, repeated TBIs are usually administered within very short intervals (hours to days), which may not allow for delayed preconditioning to develop. This may explain why this phenomenon has not been observed nor yet investigated.

Closed head injuries are often described as heterogeneous largely because of the variation in the degree of injury. Although the data in this study demonstrated that a prior mTBI consistently impaired a subsequent neurogenic or proliferative response, we noted within group variation in the degree of proliferation after the second mTBI. This variation was predicted by the degree of proliferation that occurred after the first injury (Fig. 4D). In addition, we also showed that the degree of proliferation after the first mTBI was predicted by the degree of neurologic injury (Fig. 4C). By extension, it is reasonable to expect that the NSS after the first injury should predict the proliferative response to the second injury. However, we did not find a significant correlation between these two measures (data not shown). This may have been due to a potential preconditioning effect which was most likely and apparent in the two-month injury interval group. A noteworthy caveat in this experiment is that because animals were euthanized approximately 3 weeks after BrdU administration, the density of BrdU^+^ cells reflected not only a record of cells that proliferated acutely after mTBI, but also a loss of cells that occurred in the subsequent days to weeks.

Conventionally, post-TBI neurogenesis is thought to benefit hippocampal recovery, particularly because neurogenesis in the non-injured brain plays a variety of important roles in cognitive function. Furthermore, post-TBI neurons survive and functionally integrate into the hippocampal circuitry^38,44^. Nevertheless, there is compelling evidence from the epilepsy, stroke and TBI field that aberrant neurogenesis has a maladaptive role in cognitive recovery^30,42,43,56,57^. For example, abnormal synapse formation by immature neurons after seizures may promote recurrent excitation and subsequent seizures^58,59^. Additionally, more recent studies using selective methods of ablating neurogenesis suggest post-TBI neurogenesis hinders memory outcome^42,43^; Still, there are findings suggesting that post-TBI memory is impaired by the selective ablation of neurogenesis, indicating that post-TBI neurogenesis is benficial^39^. Some of these discrepancies may be model related, particularly if the “aberrant neurogenesis”, characterized by abnormal dendritic development, is influenced by the degree of injury^57^.

Our sholl analysis suggests that in contrast to more severe TBI models, a mild concussive injury does not alter the dendritic morphology of the new neurons born after the injury. Our finding that impairments in the inability to increase neurogenesis after a second mTBI were associated with worse memory deficits further suggests that post-mTBI neurogenesis is beneficial for recovery. This result may appear to contrast another study which demonstrates that inhibiting neurogenesis after repeated mTBI improves cognitive recovery, however the interest of that study did not pertain to a loss in the neurogenic response and consisted of repeated mTBI’s with injury intervals spaced too closely together to ascertain a loss in a subsequent neurogenic response. Still, one alternative explanation for our behavioral results may involve long-term memory deficits after mTBI. We did not include the mTBI-sham group in the behavior as we did not find deficits in baseline levels of neurogenesis or proliferation at either time point, however we cannot dismiss the possibility of long-term memory deficits independent of neurogenesis. A considerable limitation of our study is the lack of a causal relationship between impairments in the neurogenic response after repeated mTBI and cognitive recovery. Future follow up studies directly manipulating neurogenesis should answer whether an impaired neurogenic response contributes to worse cognitive outcomes following repeated mTBI.

## Conclusions

Our findings demonstrate that a history of a concussive-like injury impairs the neurogenic response to a subsequent concussion and that these impairments are associated with a lack in the proliferative response of the RGCs. Importantly, we show these deficits are not due to exhaustion of the RGC pool. The different injury intervals suggest that contrary to current opinion, these post-mTBI impairments are not transient, rather are more pronounced with time. Our work also demonstrates that unlike more severe TBIs, the neurons generated after a mTBI are not aberrant in their morphology and that the inability to increase levels of neurogenesis after a repeated mTBI is associated with worse neurogenesis-sensitive memory. These results reveal a new element in the pathology of repeated TBI and provide a novel target to help maximize cognitive recovery.

## Abbreviations

DG: dentate gyrus
DCX: doublecortin
GCL: granule cell layer
ML: molecular layer
mTBI: imld traumatic brain injury
NSS: neuroseverity score
IPCs: precursor cells
RGCs: radial glia-like cells

## Acknowledgments

This material was supported by start-up funds for Dr. Villasana from the Anesthesiology and Perioperative Medicine Department at Oregon Health & Science University and by the Dow Department at Legacy Research Institute. We thank Dr. Stefanie Kaech-Petrie of the OHSU Advanced Light Microscopy Core for technical assistance with imaging; lab research assistant Sarah Feller and lab volunteer Sam Raphael for assisting with the pilot studies for this report.

